# Physical Characterization of Triolein and Implications for Its Role in Lipid Droplet Biogenesis

**DOI:** 10.1101/2021.04.21.440804

**Authors:** Siyoung Kim, Gregory A. Voth

## Abstract

Lipid droplets (LDs) are neutral lipid storing organelles surrounded by a phospholipid (PL) monolayer. At present, how LDs are formed in the endoplasmic reticulum (ER) bilayer is poorly understood. In this study, we present a revised all-atom (AA) triolein (TG) model, the main constituent of the LD core, and characterize its properties in a bilayer membrane to demonstrate the implications of its behavior in LD biogenesis. In bilayer simulations, TG resides at the surface, adopting PL-like conformations (denoted in this work as SURF-TG). Free energy sampling simulation results estimate the barrier for TG relocating from the bilayer surface to the bilayer center to be ∼2 kcal/mol in the absence of an oil lens. SURF-TG is able to modulate membrane properties by increasing PL ordering, decreasing bending modulus, and creating local negative curvature. The other neutral lipid, dioleoyl-glycerol (DAG), also reduces the membrane bending modulus and populates the negative curvature regions. A phenomenological coarse-grained (CG) model is also developed to observe larger scale SURF-TG-mediated membrane deformation. The CG simulations confirm that TG nucleates between the bilayer leaflets at a critical concentration when SURF-TG is evenly distributed. However, when one monolayer contains more SURF-TG, the membrane bends toward the other leaflet, followed by TG nucleation if a concentration is higher than the critical threshold. The central conclusion of this study is that SURF-TG is a negative curvature inducer, as well as a membrane modulator. To this end, a model is proposed in which the accumulation of SURF-TG in the luminal leaflet bends the ER bilayer toward the cytosolic side, followed by TG nucleation.

## INTRODUCTION

Lipid droplets (LDs) are energy- and lipid-storing organelles surrounded by a phospholipid (PL) monolayer.^1-4^ Enriched with neutral lipids such as triacylglycerol (TG) or sterol esters in their core, LDs are known to be formed from the endoplasmic reticulum (ER). However, little is known about LD biogenesis in spite of its implications for a number of metabolic diseases. The current model posits three distinct steps^5-7^: 1) synthesis of TG by diacylglycerol o-acyltransferase (DGAT) at the ER, 2) phase separation of TG at a critical concentration to form nascent LDs, and 3) LD budding toward the cytosol. From nucleation to budding, the lipid droplet assembly complex is known to mediate LD biogenesis. Afterwards, the complex is separated into the ER resident protein seipin and the lipid droplet assembly factor (LDAF1).^8^ The absence of this complex results in ectopic LDs; however, how this machinery catalyzes LD formation remains poorly understood.^9^

Computer simulations have clearly proven to be highly valuable for the study of biomolecules such as lipid bilayers and proteins at the scale of up to ∼100 M atoms and ∼1 milliseconds, largely enabled by rapid advances in hardware, software, enhanced sampling, and supporting theories.^10^ As such, molecular dynamics (MD) simulations have provided valuable insights on both dynamical and structural properties. For instance, the physical properties of lipid bilayer membranes have been extensively studied involving varying sizes and lipid compositions, as well as lipid-protein interactions at various resolutions (see, e.g., refs 11-15). In contrast, computational studies of LDs are only just beginning to receive attention; this is partly because the available force fields do not support certain neutral lipids. For this reason, prior computational studies have produced a new TG topology with or without a new parameter set. For instance, the Vanni group has been incorporating the Shinoda-DeVane-Klein (SDK) force field in their LD studies, which involves the parametrization of the glycerol group to reproduce interfacial tension at the water interface.^16-19^ The more common (and easier) approach is to create a new TG topology by replacing the head group of a PL with one of its tails without creating a new parameter set. This approach has been used in all-atom (AA) MD^20-23^ and coarse-grained (CG) MD simulations, the latter run with the MARTINI CG force field.^24-27^

Recent 10 µs-long AA simulations of LDs report surface-oriented TG (called SURF-TG in this work),^22^ which was also implicated in NMR experiments.^28^ The relative amount of SURF-TG to PLs, which ranges from 5-8% in the unstressed LD^22^ to ∼20% in the stressed LD,^23^ has been shown to modulate the area per lipid (APL) and interdigitation with PLs. The display of SURF-TG at the LD surface creates a unique composition and defect type that does not exist in other bilayer-bounded organelles, thereby providing a plausible explanation for LD-specific targeting or conditional targeting by certain proteins. For instance, the amount of SURF-TG increases in expanding LDs to reduce the surface tension.^23^ The highly amphipathic, autoinhibitory motif of CTP:phosphocholine cytidylyltransferase (CCT) is predicted computationally to sense defects created by SURF-TG and preferentially binds to those defects when LDs are expanding.^23^ Once fully associated, the insertion depth of the Trp278 of CCTα is aligned with the glycerol moiety of SURF-TG to form hydrogen bonds, which may increase the binding affinity.^23^ Although additional research will be needed to confirm that surface-oriented neutral lipids at the LD surface promotes specificity in LD targeting, this latter CCT binding simulation study reinforces the possible importance of SURF-TG in peptide targeting.

In this paper, the focus is not on investigating SURF-TG as a stress reducer and peptide-targeting mediator but instead on SURF-TG as a membrane property modulator. In particular, the way in which its molecular geometry is implicated in LD biogenesis is discussed. Lipids that have a conical shape such as phosphatidylethanolamine (PE), diacylglycerol (DAG), or TG do not usually form a lamellar structure by themselves.^29-31^ One notable example is 2,3-dioleoyl-D-glycero-1-phosphatidylcholine (DOPC) vs 2,3-dioleoyl-D-glycero-1-phosphatidylethanolamine (DOPE). The only difference between those two molecular structures is the head group, with DOPC having -CH_2_-N(CH_3_)_3_+ in comparison to the -CH_2_-NH_3_+ head group of DOPE. Therefore, a single DOPC molecule features a cylindrical shape, while DOPE has a conical shape. This difference in molecular topology leads to two distinct phases when DOPC and DOPE exist in single-component bulk systems: DOPC forms a planar bilayer while DOPE mostly forms an inverse hexagonal phase.^32-33^ The physical properties of these two phases have been characterized experimentally^32, 34-36^ and computationally.^37^ Compared to DOPC or DOPE, DAG is more conical because of the absence of the head group, and SURF-TG is even more so due to the addition of the acyl chain. Using AA-MD simulations, we demonstrate here that these conical lipids are able to modulate membrane properties, such as bending modulus, and populate negative curvature regions of the membrane.

The present study also describes the development of a phenomenological CG model for PL and TG to access larger effective simulation time scales, wherein we show that TG nucleates at a critical TG concentration between the leaflets as opposed to the asymmetric distribution of SURF-TG, which induces membrane bending first. Collectively, we demonstrate that SURF-TG is a negative curvature inducer. To this end, we propose a model in which SURF-TG-driven bending helps budding and determines the directionality of LD budding.

## METHODS

### AA-MD simulations and system setup

In this study, dioleoyl-glycerol (DOGL) and triolein were used to represent DAG and TG, respectively. A system was prepared using either the CHARMM-GUI membrane builder^38-41^ or PACKMOL.^42^ The details for each simulation and system are summarized in Table 1. The TG molecular structure used for the PACKMOL input was based on a prior LD simulation featuring a trident structure.^22^ Three different TG models were used (Fig. 1). The first one was made based on the CHARMM36 (C36) lipid force field^43^ by replacing the head group of DOPE with its *sn-1* acyl chain. However, the structural and thermodynamic properties of nonpolar molecules are sensitive to the cutoff distance of Lennard-Jones (LJ) interactions, which was discussed in ref ^44^. Indeed, the bulk TG simulations, using the 1.2 nm cutoff distance of LJ interactions, which is consistent with C36, resulted in the low density and low surface tension at the vacuum interface (See Fig. 2 in the Results). Recently, a LJ cutoff-free version of C36 (C36/LJ-PME) was released.^45-46^ The second TG model was made based on the C36/LJ-PME parameters in the same fashion used in the first model. However, both first and second models suffer from the low surface tension at the water interface (Fig. 2 in the Results). Therefore, we introduce the revised TG model (C36/LJ-PME-r), which was made based on C36/LJ-PME, however, with the reduced partial charge distribution at the ester group (Fig. 1). The new partial charge distribution was determined in a manner to match the surface tension of TG at the water interface. The parameters of the ester group were originally derived to best describe the PL behavior at the water interface; therefore, we hypothesized that the partial charges of the TG ester group should be reduced, which is a non-polar molecule. The suggested charge distribution (Fig. 1) was obtained by conducting a series of tests: Adding Δq to the oxygen atoms (O11, O12, O21, O22, O31, and O32) and subtracting 2Δq from the carbon atom (C11, C21, and C31). When Δq = 0 (C36/LJ-PME), Δq = 0.05, Δq = 0.10, and Δq = 0.20 (C36/LJ-PME-r), the surface tension was 13.7 mN/m, 16.4 mN/m, 24.9 mN/m, and 32.2 mN/m, respectively. A systematic way to optimize parameters, not only partial charges but also LJ parameters, will be a focus of future work (see, e.g., ref^46^ for an example). The partial charges of PL were unchanged. Therefore, for the simulations that do not include TG, C36/LJ-PME and C36/LJ-PME-r are the same. All of those TG topologies are available in https://github.com/ksy141/TG. Of note, DAG is also not included in the original C36 force field release; nonetheless, the CHARMM-GUI developers (The Im group) created a new topology for DAG based on the existing parameter set.

**Table 1.** Description of AA simulations.

| AA Simulations | Lipid composition <sup>a)</sup> | Time | Ensemble | Replicas | Setup |
| --- | --- | --- | --- | --- | --- |
| Bulk TG | 216 TG | 100 ns | NPT | 6 <sup>b)</sup> | PACKMOL |
| Bulk TG-vacuum | 216 TG | 200 ns | NVT | 6 <sup>b)</sup> | From above sim. |
| POPC bilayer | 128 POPC | 200 ns | NPT | 1 | PACKMOL |
| POPC + TG bilayer | 128 POPC + 1 TG | 1 μs | NPT | 1 | PACKMOL |
| POPC + TG bilayer | 128 POPC + 1 TG | 300 ns | NPT + biased | 12 <sup>c)</sup> | From above sim. |
| POPC + TG bilayer | 128 POPC + 1 TG | 2 μs | NPT + biased | 5 | From above sim. |
| POPC + TG bilayer | 124 POPC + 8 TG | 1 μs | NPT | 1 | PACKMOL |
| POPC + TG bilayer | 1984 POPC + 128 TG | 600 ns | NPT | 1 | From above sim. |
| POPC bilayer | 4050 POPC | 300 ns | NPT | 1 | CHARMM-GUI |
| POPC + DAG bilayer<br>(referred to as DAG 10%) | 3200 POPC + 320 DAG | 1 μs | NPT | 1 | CHARMM-GUI |
| POPC + DAG bilayer<br>(referred to as DAG 20%) | 3200 POPC + 640 DAG | 1 μs | NPT | 1 | CHARMM-GUI |
| POPC + DAG bilayer<br>(referred to as DAG 30%) | 3200 POPC + 960 DAG | 1 μs | NPT | 1 | CHARMM-GUI |
| DAG 30% membrane<br>deformation and then NPAT | 3200 POPC + 960 DAG | 20 ns +<br>1 μs | NPγT and<br>then NPAT | 1 | From above sim. |
| DOPC bilayer | 128 DOPC | 200 ns | NPT | 1 | CHARMM-GUI |

a) The total number of lipid molecules of each type in a system.
b) With varying LJ cutoff distances
c) 12 windows

**Figure 1.**
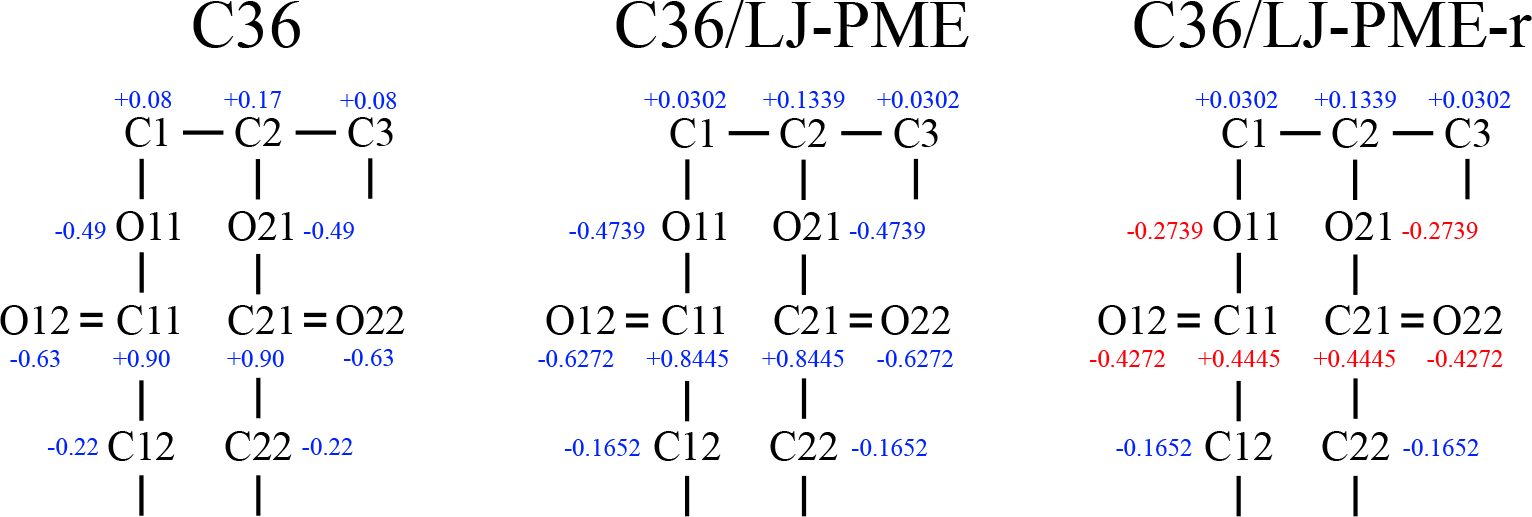
The glycerol moiety of three TG models with partial charges. The revised TG model (C36/LJ-PME-r) has a significantly reduced charge distribution in the ester group (shown in red).

**Figure 2.**
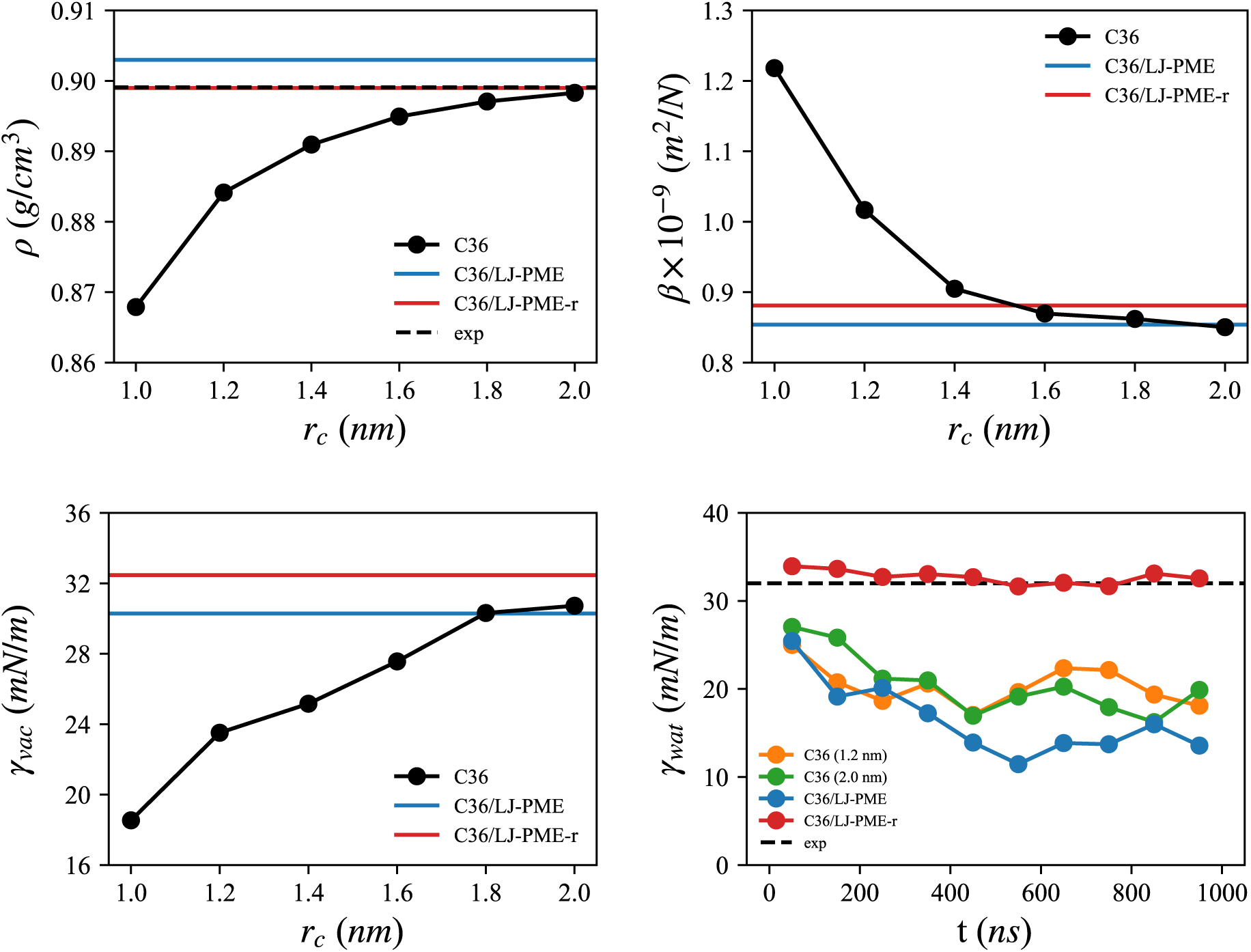
Physical properties of the bulk TG evaluated with three different TG models. For C36, the properties were calculated as a function of LJ cutoff distances and the force switching function was applied between *r*_*c*_ - 0.2 *nm* and *r*_*c*_. For the surface tension at the water interface, the values were averaged every 100 ns. C36 (1.2 nm) represent the simulation carried out with C36 using the cutoff distance of 1.2 nm.

All simulations were run by GROMACS 2018 and 2020^47^ with the C36^43^, C36/LJ-PME ^45-46^, or C36/LJ-PME-r force field. Simulations using C36 used the 1.2 nm LJ cutoff distance with a force-switching function between 1.0 nm to 1.2 nm, unless otherwise noted. All simulations including DAG were performed with C36. Simulations using C36/LJ-PME and C36/LJ-PME-r used the real-space cutoff distance of 1.0 nm. Biased simulations (described below) were performed with the external plugin, PLUMED2 (v.2.6).^48^ Simulations were evolved with a 2-fs timestep. The long-range electrostatic interactions were evaluated with the Particle Mesh Ewald algorithm,^49^ with the distance cutoff set to be the same as the LJ interaction. Every bond involving a hydrogen atom was constrained using the LINCS algorithm.^50^ The Nose-Hoover thermostat was used to maintain a temperature of 300 K (simulations involving DAG) or 310 K with a coupling time constant of 1 ps.^51-52^ For the constant pressure simulations, the Parrinello-Rahman barostat was used to maintain a pressure of 1 bar with a compressibility of 4.5 × 10^−5^ bar^-1^ and a coupling time constant of 5 ps.^53^ For membrane simulations, the pressure was coupled semi-isotropically, whereas during bulk simulations the pressure was coupled isotropically. For constant area simulations (NPAT), only the pressure of the Z dimension was controlled. In order to deform the DAG 30% membrane, a constant surface tension simulation (NPγT) was conducted for 20 ns with the Berendsen thermostat and barostat using the same parameters.^54^ The pressure was coupled semi-isotropically with the pressure in the X and Y dimension set to −100 bar during membrane deformation.

### Density, isothermal compressibility, and surface tension

Density was obtained with the GROMACS analysis tool, which was outputted every 2 ps. Isothermal compressibility was evaluated from

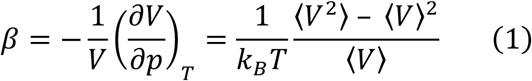

where the bracket represents the ensemble average and V denotes the volume of the system. Similarly, GROMACS was used to analyze the surface tension at the vacuum and water interfaces. Bilayer area compressibility was determined using the following equation

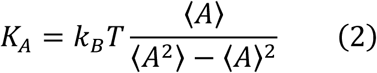

where *A* is the instantaneous total area of a membrane.^55^

### Order parameters

Order parameters were evaluated from *S*_*CD*_ = 0.5 × |〈3 cos^2^ *θ* - 1〉|, where the angle (*θ*) is between the Z axis and the position vector of an acyl carbon atom to a bonded hydrogen atom; the bracketed portion indicates the ensemble average.^56-58^

### Potential of mean force of TG flip-flop

In order to estimate the preferential location of TG in a bilayer and its tendency to become SURF-TG, the potential of mean force (PMF) of the TG as a function of its Z position in the bilayer was calculated. A 3-palmitoyl-2-oleoyl-D-glycero-1-Phosphatidylcholine (POPC) bilayer membrane consisting of 128 POPC molecules and one TG molecule was prepared. Biased simulations were performed with the collective variable representing the Z distance between the center of the mass of phosphorus atoms and the center of the mass of the glycerol moiety of the TG molecule. Replica-exchange^59^ umbrella sampling^60^ (REUS) simulations were run for 300 ns with a harmonic restraint of a force constant of 47.8 kcal/mol/nm^2^ for each window. The total of 12 windows was prepared with a 0.2 nm spacing over a range of 0.0 to 2.2 nm (0 is the average Z position of the center of the mass of the phosphorus atoms, which is comparable to the center of the membrane). An initial structure for each window was prepared with a steered MD simulation. The exchange between windows was attempted every 2,000 steps. The weighted histogram analysis method (WHAM) was used for the calculation of the PMF with a bin spacing of 0.0275 nm.^61^ The Grossfield’s WHAM implementation was used.^62^ The simulations were divided into five equal-length blocks to estimate the errors. The PMF was calculated for each block and then the standard deviation of block PMFs was reported as the errors. To support the PMF result, we also carried out efficient, fast-converging transition-tempered metadynamics (TT-MetaD) simulations with the same collective variable.^63-64^ The Gaussian hill was deposited every 5K steps at a height of 0.29 kcal/mol and width of 0.08 nm or 0.1 nm. The bias factor of 6 and threshold of 2.4 kcal/mol for TT-MetaD were used. The position of the two basins were set to −1.1 nm and +1.1 nm, when the center of the mass of phosphorus was set to zero. Five simulation replicas, each 2 µs long, were performed to estimate the averaged potential of mean force (PMF). The errors represent the standard deviation of the PMFs obtained from the five replicas.

### CG simulations

In order to study SURF-TG-driven membrane deformation and nucleation, a new phenomenological implicit solvent CG model was developed for the PL and TG. Each PL and TG molecule consisted of 11 and 13 CG beads, respectively. Each acyl chain was composed of 4 CG beads, with 2 different CG types. The approximate molecular groupings of the PL and TG structures are shown in Fig. S1. In this model, the attraction was modeled with the Gaussian function, -*A exp*(-*Br*^2^) where *B* = 2 *nm*^−2^, and the repulsion with the repulsive LJ interaction, 4*ε*(*σ/r*)^12^ where ξ = 0.0028 *kcal*/*mol*. No CG bead carried any charge. The attraction only occurs between hydrophobic CG beads (beads in the acyl chain or CT1 and CT2 in Fig. S1 of the Supporting Information), between the glycerol CG beads of TG, and between the glycerol CG beads of TG and PL (No attraction exists between the glycerol CG beads of PL); in contrast, repulsion occurs between all CG beads unless the pair is bonded. The attraction and repulsion parameters, *A* and *σ*, were primarily parameterized to reproduce the radial distribution function (RDF) from the mapped AA (C36) RDF (Figs. S2 and S3). Unless otherwise indicated, the repulsion parameter (*σ*) of a pair composed of two different types of CG beads was determined using an arithmetic average, while the corresponding attraction parameter (*A*) for such a pair was set explicitly. All CG bonds were modeled with the harmonic potential, with the equilibrium distance of 0.5 nm and the force constant of 500 kcal/mol/nm^2^. The angle potentials were modeled with the harmonic potential with a small force constant of 0.12 kcal/mol/rad^2^. However, we note that the properties of PL and TG in general were not sensitive to the bond and angle force constants. The details of this CG model can be found in Fig. S1. The potentials and inputs are available at https://github.com/ksy141/SK_CGFF.git.

All CG simulations were conducted with GROMACS 2018 using a 20-fs timestep.^47^ The cutoff distance of 2.4 nm was used for non-bonded interactions. Temperature was controlled via velocity rescaling with a stochastic term with the target temperature of 310 K and a time constant of 1 ps.^65^ A pressure of 1 bar was maintained with the Parrinello-Rahman barostat with a compressibility of 4.5 × 10^−5^ bar^-1^ and a coupling time constant of 10 ps. For the PL or PL+TG bilayer simulation, the pressure was semi-isotopically coupled with the X and Y dimensions but was not coupled with the Z dimension (by setting compressibility in the Z dimension to 0). For the bulk TG simulation, the pressure was isotopically coupled. For bilayer simulations that have asymmetric distribution of TG between two leaflets, weak pressure and temperature couplings were used to minimize perturbations of the evolution of the membrane remodeling. For those simulations that convey membrane deformation, time constants of 1 ns and 10 ns for temperature and pressure were used, respectively. Note that a previously reported implicit solvent CG model that detected membrane deformation used weak couplings as well.^66^ Finally, the TT-MetaD simulations were carried out to obtain the TG flip-flop PMF. The Z position of the TGL atom respect to the membrane center was biased with the same TT-MetaD parameters used in the AA simulations. The initial structures were prepared using PACKMOL^42^ and MDAnalysis^67^; simulation details are provided in Table 2.

**Table 2.**
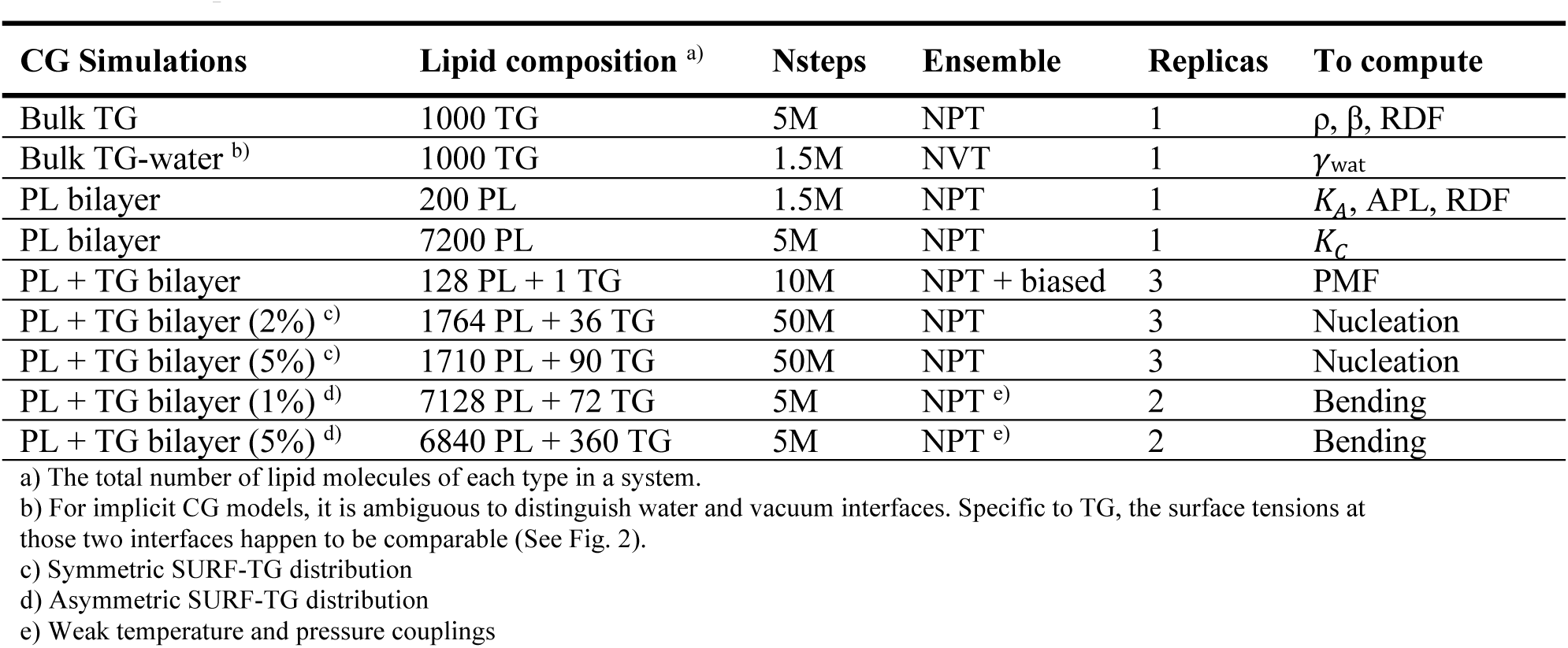
Description of CG simulations.

### SDK CG simulations

To add additional support to the findings and conclusions, CG simulations with the SDK force field (SPICA force field)^68-70^ were carried out with the LAMMPS MD package.^71^ Two systems were prepared: One TG molecule in a bilayer containing 128 POPC molecules to obtain the TG flip-flop PMF and a POPC + DOPE bilayer immersed in water (or a nanodisc) to show bending. The TG force field that reproduces the surface tension at the water interface was used in the first system.^16^ In the second system, asymmetric distribution of PLs was used with the upper leaflet having 576 POPC molecules and the lower leaflet having 522 POPC and 54 DOPE molecules. Simulations were evolved with a 10-fs timestep. The Nose-Hoover thermostat^51^ and barostat^53, 72-73^ were used to maintain a temperature of 310 K (first system) or 300 K (second system) and a pressure of 1 bar with damping parameters of 20 ps and 50 ps, respectively. The cutoff distance of 1.5 nm was used, and the long-range electrostatic interactions were evaluated with the particle-particle-particle-mesh (PPPM) solver. The force error of 10^−5^ kcal/mol/Å, the third order, and the grid size of 20 Å in each dimension were used for the long-range electrostatic interactions.

### Analysis and visualization

Simulations were analyzed using MDAnalysis^67^ and GROMACS.^47^ Unless otherwise noted, the standard errors (*se*) were reported in this work. Equilibrated trajectories were first divided into *M* blocks of equal length and the block average for each block was calculated. The standard errors (*se*) were estimated by 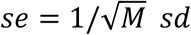, where *sd* is the standard deviation of the block averages. Five blocks (*M* = 5) were used in this work. Molecular images were captured using PyMOL.

## RESULTS

### AA-MD

#### Physical properties of bulk TG

In this section, we establish the quality of three TG models (C36, C36/LJ-PME, and C36/LJ-PME-r; see Fig. 1 and Methods) by characterizing four physical properties of bulk TG (Fig. 2): density (ρ), isothermal compressibility (β), surface tension at the vacuum (γ_vac_), and surface tension at the water interface (γ_wat_). With a prior study demonstrating that the first three properties are sensitive to the LJ cutoff distances,^44^ we performed MD simulations (C36) of a bulk TG consisting of 216 TG molecules with varying LJ cutoff distances. The computed density and isothermal compressibility of the bulk TG were shown to converge when the LJ cutoff distance was increased from 1.0 nm to 2.0 nm (Fig. 2). The density of the bulk TG approached the experimental value (0.8991 g/cm^3^ at 313 K)^74^ when simulated with the LJ cutoff of 2.0 nm (0.898 g/cm^3^). The same calculations were carried out with C36/LJ-PME and C36/LJ-PME-r. The extended LJ interactions caused a higher density in the C36/LJ-PME results, however, the extra increase became compensated by the reduced charge distribution in C36/LJ-PME-r. The same behavior was observed in the RDF of the TG glycerol moiety (Fig. S4). While C36/LJ-PME increases the first peak due to the increased LJ range, the reduced charge distribution in C36/LJ-PME-r lowers the increased peak. Next, we performed MD simulations of the TG-vacuum interface by adding an empty space in the Z dimension to the bulk TG system. Consistent with density and isothermal compressibility, the surface tension of the TG-vacuum interface converges with the increased LJ cutoff distance and also agrees reasonably well with the results of C36/LJ-PME and C36/LJ-PME-r. In all cases, no TG molecules were released from the bulk TG. Although there are no experimental TG data for β and γ_vac_, the results of C36 with the 2.0 nm cutoff distance, C36/LJ-PME, and C36/LJ-PME-r show the reasonable agreement for ρ, β, and γ_vac_.

On the other hand, the experimental surface tension at the water interface (32 mN/m)^75^ was not reproduced except for C36/LJ-PME-r (Fig. 2). The surface tension became reduced to < 20 mN/m when using C36 or C36/LJ-PME-r, verifying the necessity of reducing the distribution of partial charges in the ester group. Although the additive force field cannot take into account polarization therefore cannot accurately produce the water density in nonpolar environment unless further modified,^76-77^ it is worth nothing the large difference found in the water density at the TG layer. While C36 and C36/LJ-PME demonstrate the high water density (∼10^−2^ g/cm^3^), C36/LJ-PME-r shows the low water density (∼10^−3^ g/cm^3^), which agrees with the experimentally measured density (1.8 × 10^−3^ g/cm^3^).^78^

#### TG flip-flop

We characterized the physical properties of TG in a bilayer membrane in order to investigate its implication in LD biogenesis. A bilayer membrane containing 128 POPC molecules and 1 TG molecule was prepared. We then calculated the PMF of the TG molecule (Fig. 3) as a function of its Z position within a bilayer membrane by performing the REUS simulations. Both C36 and C36/LJ-PME-r demonstrate the stability of SURF-TG in the absence of an oil lens although the C36 result overestimates the stability at the surface by ∼1.5 kcal/mol compared to the C36/LJPME-r result. The preferential location of the TG molecule is slightly different by 0.2 nm as well. We further verified our results with another biased methodology. Five replicas of TT-MetaD simulations were carried out with the same collective variable, each run for 2 µs. The PMFs obtained from the REUS and TT-MetaD simulations show good agreement (Fig. S5). Finally, the 1-µs unbiased MD simulation behaves as consistent with our findings. For the C36 simulation, the TG molecule was initially located at the center of the bilayer (CORE-TG), however, became SURF-TG rapidly (< 10 ns). During 1 µs, the TG molecule did not flip-flop but resided at the same leaflet. In contrast, the TG molecule visited the membrane center more frequently and eventually flip-flop at ∼820 ns with C36/LJ-PME-r. Taken together, both unbiased and biased results suggest that a single TG molecule is principally SURF-TG in a bilayer membrane, adopting the PL-like conformation, in the absence of an oil lens.

**Figure 3.**
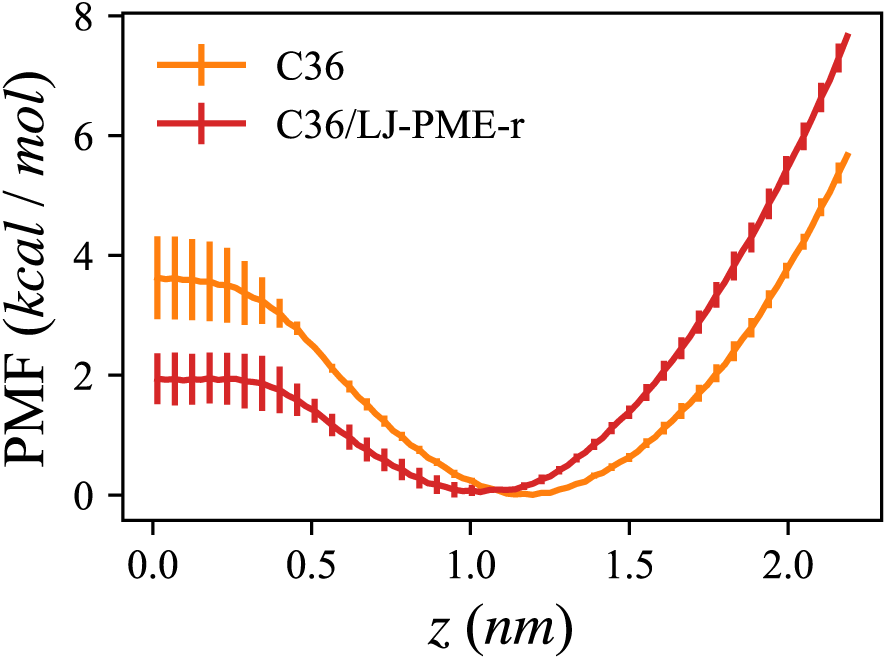
PMF of a TG molecule in a POPC bilayer as a function of the Z distance from the bilayer center (TG flip-flop), calculated from the REUS simulations. The Z position of the center of the phosphorus atoms was zeroed. The errors represent the standard deviation of the PMFs obtained from the five equal-length blocks.

#### Local membrane deformation by SURF-TG

Conical molecules such as PE or DAG are known to be responsible for amphipathic peptide targeting by creating packing defects.^79-83^ However, the precise mechanisms for how those molecules modulate membrane properties are poorly understood. The molecular shape of SURF-TG is inevitably conical, with the glycerol moiety forming the vertex of the cone and the end group of three acyl chains forming a flat base while POPC features a quasi-cylindrical shape. In order to study how SURF-TG modulates the local properties of bilayers, we computed the height field of the phosphorus atoms of the upper leaflet in the bilayer containing 128 POPC molecules and 1 TG molecule. Histogram analysis was used for the 1-µs C36/LJ-PME-r simulation to investigate the relative positions of the phosphorus atoms of the upper leaflet with respect to the glycerol moiety of the SURF-TG molecule (Fig. 4). Approximately one-fifth of the frames were not used for this analysis because in those frames SURF-TG transitioned to CORE-TG or resided at the other leaflet (flip-flop occurred at ∼820 ns). The height of the phosphorus atoms near the SURF-TG molecule can be as low as ∼0.15 nm compared to those of the other phosphorus atoms (Fig. 4). The range of the local deformation created by one SURF-TG molecule can be more than 2 nm in both the X and Y dimensions (Fig. 4). We anticipate that if SURF-TG molecules cluster at the surface there will be stronger and longer PL packing discontinuity. We confirmed that the same conclusion can be made with the C36 trajectory (data not shown). Our data indicate that SURF-TG locally pulls PLs toward the bilayer center, thereby creating local negative curvature. How this process serves to remodel bilayer membranes will be discussed later via our CG simulation results.

**Figure 4.**
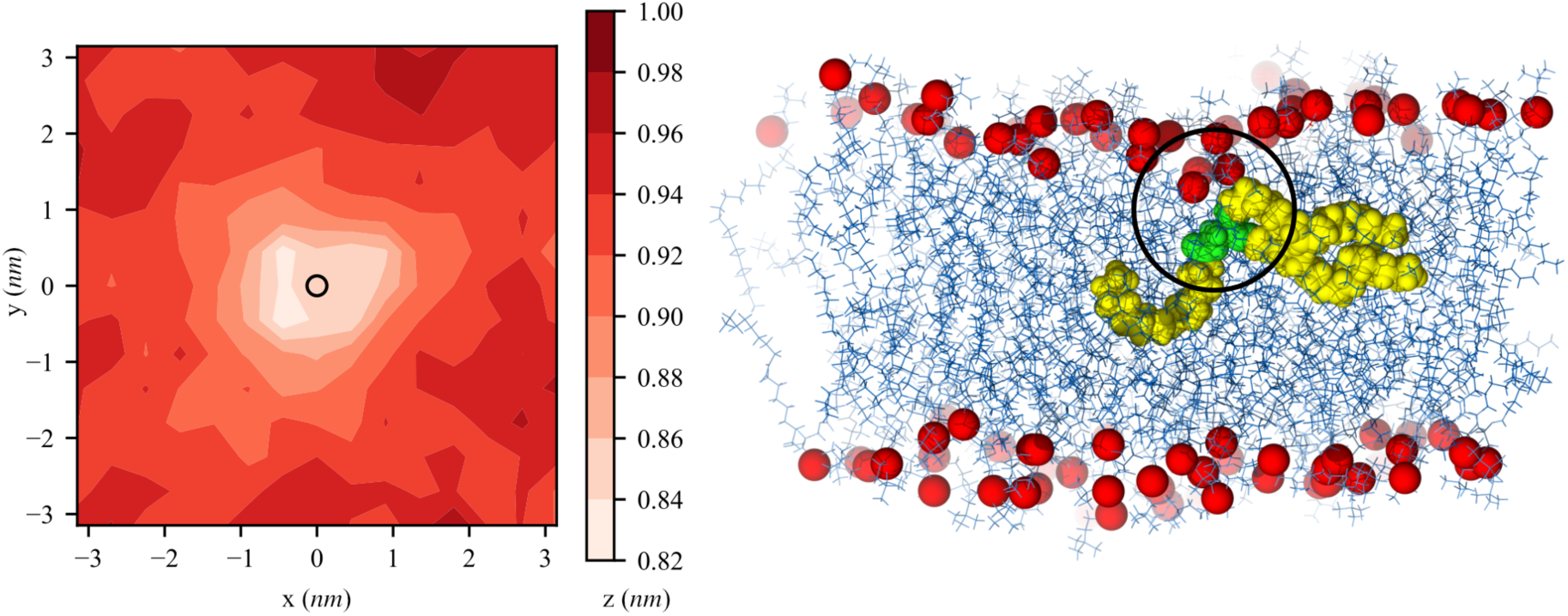
*(left)* Height field, z(x, y), of the phosphorus atoms of the upper leaflet respective to the glycerol moiety of the SURF-TG molecule (circle at the origin). Frames in which SURF-TG resided at the upper leaflet were used for the analysis. *(right)* Snapshot demonstrating the local deformation by SURF-TG. The red spheres are the phosphorus atoms and PLs are shown in blue lines. The glycerol moiety and acyl chains of TG are shown with green and yellow spheres, respectively. The simulation was carried out with C36/LJ-PME-r.

#### Molecular and hydration properties of SURF-TG

In order to characterize the molecular and hydration properties of SURF-TG, a new POPC bilayer membrane containing 6% TG (62 POPC and 4 TG molecules per leaflet) was prepared and compared with a pure POPC bilayer membrane (64 POPC molecules per leaflet). The system contained a higher concentration of TG than the critical nucleation concentration, which was reported to be 2.8% g TG or 2.4% mol TG.^28^ However, within the AA simulation timescale, TG nucleation did not occur, and all the TG molecules remained SURF-TG most of the times. The simulations were carried out with C36/LJ-PME-r. For a pure POPC bilayer membrane, C36/LJ-PME and C36/LJ-PME-r are equivalent.

The order parameters of SURF-TG and POPC were computed (Fig. 5). Interestingly, the order parameters of POPC in the POPC+TG bilayer (continuous lines) were found to be larger than those of POPC in the pure POPC bilayer (dashed lines), suggesting that SURF-TG increases the order of POPC molecules. As reported in a previous study, the increase in PL ordering in LDs compared to the bilayers can be attributed to the fact that CORE-TG interdigitates with the PL monolayer and increases the density of the low-density tail region.^22^ Similarly, short LD simulations lacking any SURF-TG also demonstrated increased PL ordering.^84^ Our results suggest that SURF-TG increased PL ordering as well. The SURF-TG order parameters have the same trend with PL; however, they are more reduced than PL. The reduced order parameters of TG compared to PL can be attributable to the higher degree of freedom in the Z dimension of TG molecules. While PL moves little in the Z dimension and therefore its order parameters are usually determined by the lipid-packing density in the XY dimensions, TG is relatively free in moving in the Z dimension.

**Figure 5.**
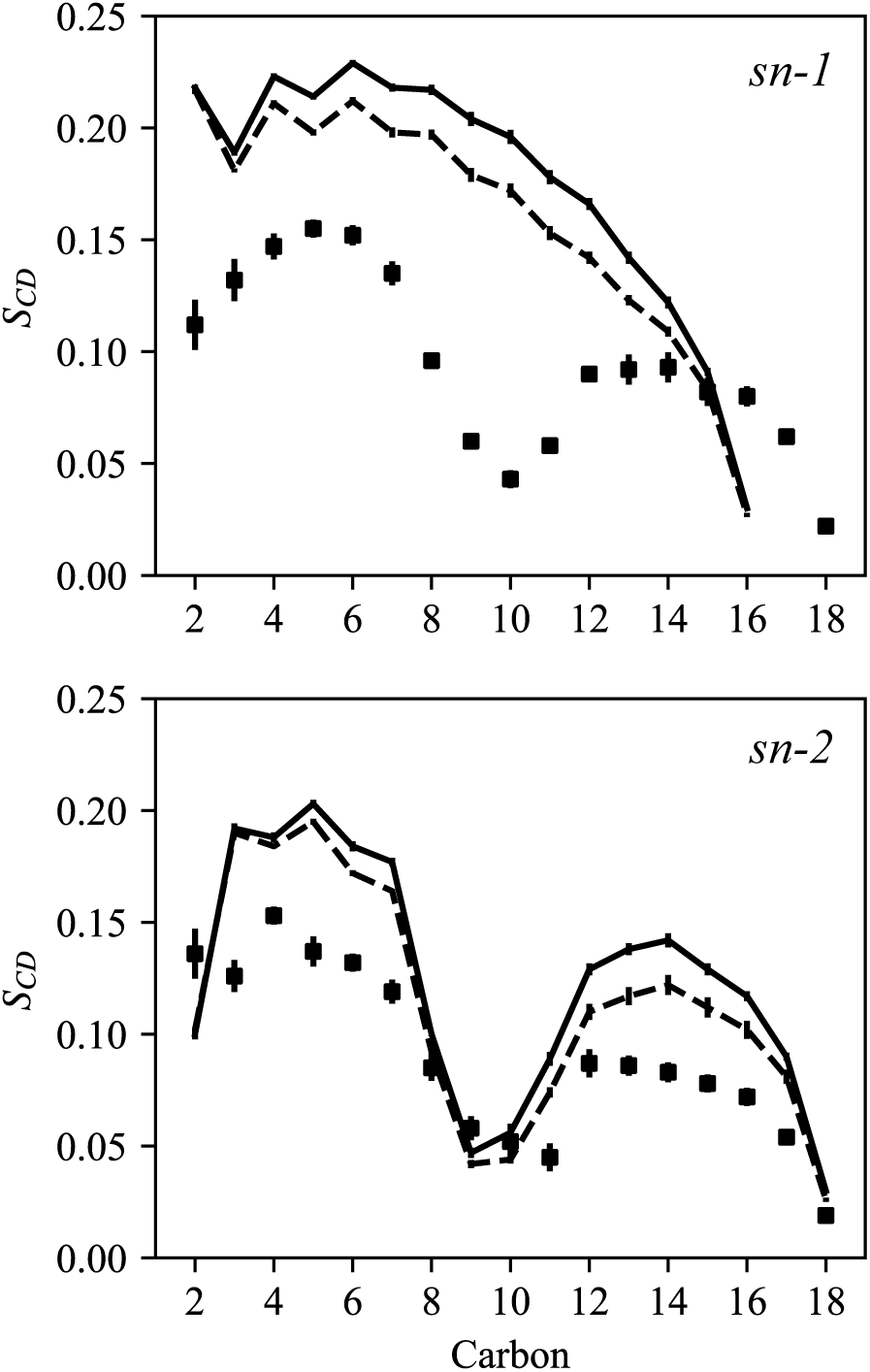
Order parameters of *sn-1* (top) and *sn-2* (bottom) chains. The POPC order parameters of the pure POPC bilayer (served as a reference) are shown in the dashed lines. The POPC order parameters of the POPC + TG bilayer are indicated in the continuous lines. The TG order parameters of the same system are shown in squares. The *sn-3* chain of TG is not included for visual clarity. The simulations were carried out with C36/LJ-PME-r.

In the same system, we characterized the orientation of each TG molecule by calculating the angle between the Z-axis and the positional vector of the center of the mass of the TG glycerol moiety from the center of the mass of the TG acyl chains. The angle as a function of the Z position is shown in Figure S6. Consistent with the TG flip-flop PMF (Figure 3), TG molecules principally reside at the membrane surface (z = ∼1.2 nm). Also, from the orientation analysis, we were able to confirm that TG adopts PL-like conformation in which the glycerol moiety is exposed to the membrane surface and its acyl chains are extended toward the membrane center.

The hydration properties of SURF-TG were evaluated in the same POPC+TG bilayer. It was suggested that the SURF-TG has the primary carbonyls in *sn-1* and *sn-3* chains that are more exposed to water than the secondary carbonyl in *sn-2* chain, which would explain specificity for hydrolysis at the primary carbonyl position.^28^ The computed RDF between each TG oxygen atom (O11, O21, and O31) and water shows reduced accessibility of water to the secondary carbonyl (Fig. 6). However, the RDFs between the other oxygen atoms (O12, O22, O32) and water were identical at the first peak (data not shown).

**Figure 6.**
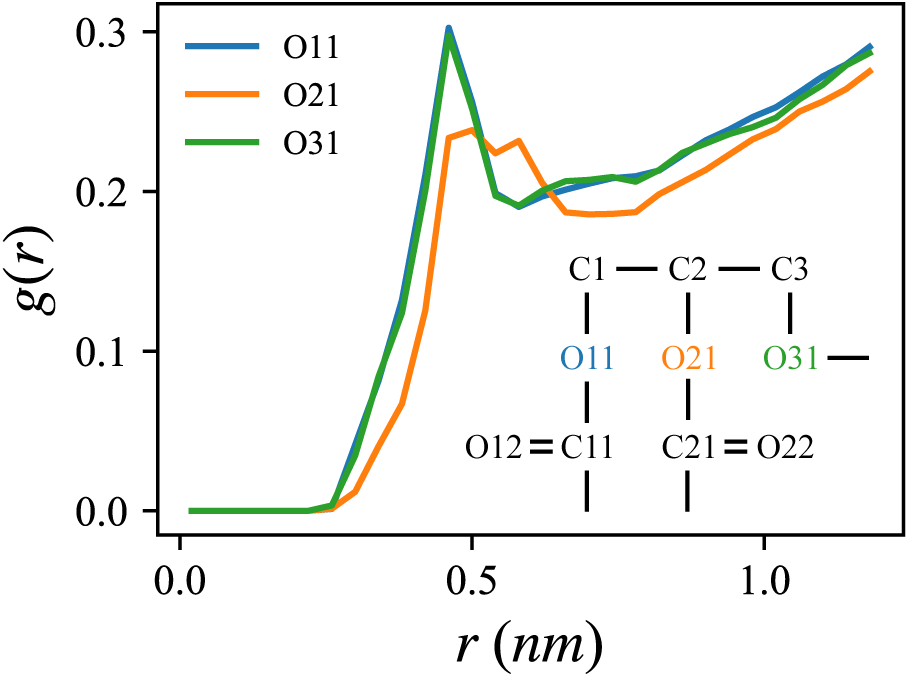
The RDFs of TG oxygen atoms (O11, O21, O31) with the oxygen atom of water. The structure of the TG glycerol moiety is illustrated in the inlet. The simulation was carried out with C36/LJ-PME-r.

#### Impact of neutral lipids on membrane properties

In order to systematically study the impact of neutral lipids on bending modulus (*k*_*c*_), we assembled large POPC bilayer membranes of varying DAG concentrations ranging from 0% to 30% and computed the bending modulus using the undulation spectrum.^85-87^ On our simulation timescale (1 µs), there was no evidence for phase separation in the DAG, despite its high concentration. With an increased concentration of DAG, the bending modulus decreased (Fig. 7), suggesting that DAG lowers bending rigidity. This agrees with the previous simulation showing the decreased bending modulus of a bilayer membrane containing DAG.^88^

**Figure 7.**
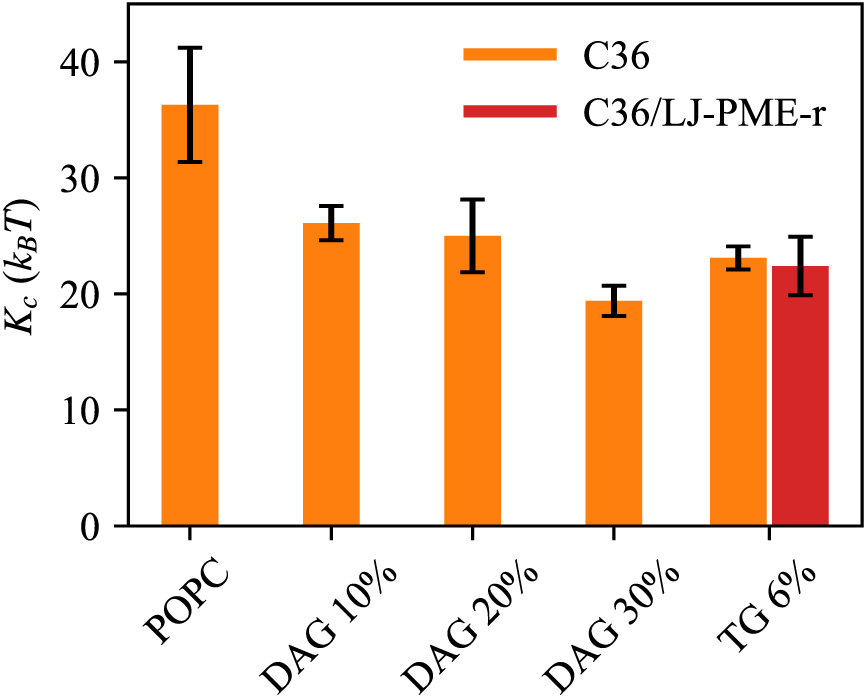
Bending modulus of five POPC bilayer membranes, containing different amounts of DAG or TG. The number of POPC and DAG or TG molecules in each system is as follows: POPC (POPC 4050), DAG 10% (POPC 3200 + DAG 320), DAG 20% (POPC 3200 + DAG 640), DAG 30% (POPC3200 + DAG 960), TG 6% (POPC 1984 + TG 128). The errors represent the standard errors.

We also confirmed a similar correlation for the bilayer containing 6% SURF-TG with both C36 and C36/LJ-PME-r (Fig. 7). The bending modulus of this particular bilayer was determined to be 23.1 ± 1.0 *k*_*B*_*T* (C36) and 22.4 ± 2.5 *k*_*B*_*T* (C36/LJ-PME-r), which is smaller than 32 *k*_B_*T* (ref 12) or 36 ± 5 *k*_*B*_*T* (Fig. 7), the bending modulus of the pure POPC bilayer. The results are supported by the recent experiments showing the reduced bending modulus of a bilayer membrane containing TG.^89^

Next, constant surface tension was applied semi-isotropically for 20 ns to induce deformation of the 30% DAG membrane. The resulting structure displayed a buckling that resembled a cubic function in the X axis (Fig. 8a). We then ran constant area simulation (NPAT) for 1 µs, confirming that the overall buckling was maintained (Fig. 8a and 8b). As shown in Fig. 8b, the height field region in the upper leaflet where X is between 5 nm and 15 nm displayed a negative curvature; conversely, the region where X is between 25 nm and 30 nm displayed a positive curvature. In contrast, the curvature becomes inverted in the lower leaflet (Fig. S7). Consistent with our expectations, a conical DAG is less populated at the positive curvature (for instance, the region where X is between 25 nm and 30 nm in the upper leaflet), but more populated at the negative curvature (Fig. 8c and Fig. S7). The central conclusion derived from these simulations is that a neutral lipid reduces the bilayer bending modulus and behaves as a negative curvature inducer.

**Figure 8.**
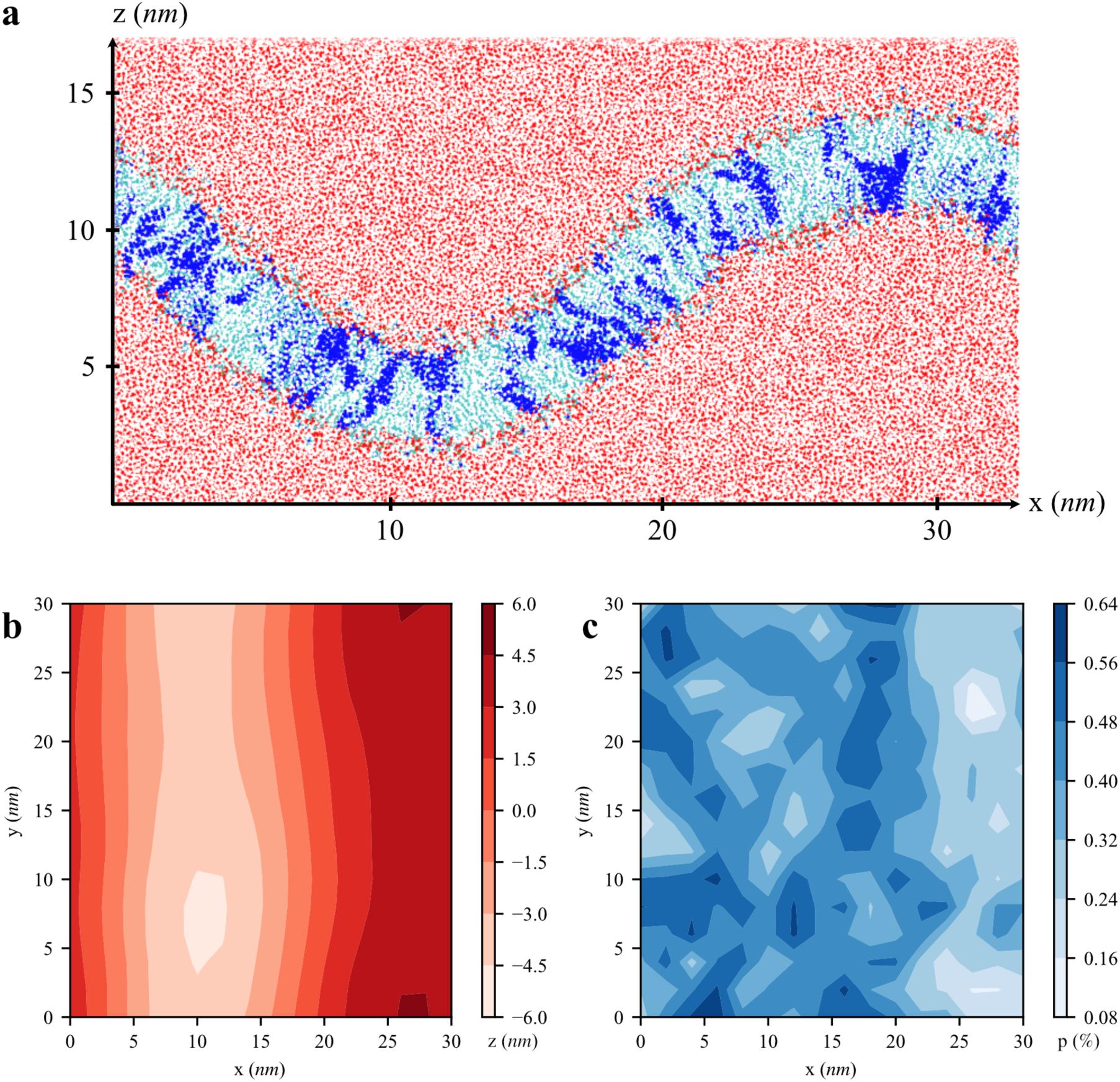
Buckled POPC+DAG membrane (DAG 30%). a) The snapshot of the MD simulation. Water is in red and DAG in blue. b) The height field, z(*x, y*), of the upper leaflet, averaged over the trajectory. The averaged Z position of the upper phosphorous atoms was zeroed. c) The DAG distribution, *p* (*x, y*), in the upper leaflet, averaged over the trajectory.

### CG-MD

#### TG bulk and PL bilayer

In order to further investigate the properties of SURF-TG in a bilayer membrane and establish their implications in LD biogenesis, we developed a new implicit solvent CG model for PL and TG as noted earlier. In this model, each PL and TG molecule consisted of 11 and 13 CG atoms, respectively, where each acyl chain was composed of 4 CG atoms and of 2 CG types. No charges were associated with CG beads. The interactions were modeled with a Gaussian function for attraction, and the LJ repulsive potential for repulsion. The details of the CG model can be found in Fig. S1.

The attraction and repulsion parameters were then primarily parameterized to reproduce the RDFs from the AA trajectories with the atoms mapped on the CG sites of the CG model (the so-called “mapped AA model”). A PL bilayer simulation consisting of 200 residues was performed for 1.5M steps, after which the computed 2-dimensional RDFs were compared with those from the mapped AA trajectory provided in Fig. S2. Similarly, a bulk TG simulation consisting of 1,000 residues was conducted for 5M steps; the computed RDFs from this trajectory were compared with those from the mapped AA trajectory shown in Fig. S3. The results showed reasonable agreement between the CG RDFs and the mapped AA RDFs, which were achieved both in the PL bilayer and bulk TG systems. We then determined the physical properties of the bulk TG and PL bilayer from the same simulations as shown in Table 3. Although the isothermal compressibility of the bulk TG obtained from the CG simulation was found to be a factor of three larger, and the bending modulus of the PL bilayer was recorded to be 1.4 times higher than the AA results, the remaining properties agreed well with the experimental or AA data.

**Table 3.** Physical properties of DOPC bilayer and TG bulk from experiments, AA simulations, and CG simulations. Standard errors are given for the simulations that we performed.

|  | Exp. | AA | CG |
| --- | --- | --- | --- |
| DOPC bilayer |  |  |  |
| APL [$nm^2$] | 0.674<sup>a)</sup>, 0.724<sup>b)</sup> | $0.70 \pm 0.00$ | $0.72 \pm 0.00$ |
| $K_A$ [$mN/m$] | 300<sup>c)</sup> | $251 \pm 20$, 290<sup>d)</sup> | $307 \pm 9$ |
| $K_C$ [$k_B T$] | 21.2<sup>e)</sup> | 29<sup>d)</sup> | $41 \pm 1$ |
| TG bulk |  |  |  |
| $\rho$ [$g/cm^3$] | 0.8991<sup>f)</sup> | $0.90 \pm 0.00$<sup>g)</sup> | $0.90 \pm 0.00$ |
| $\beta \times 10^{-9} [m^2/N]$ | N/A | $0.88 \pm 0.01^g)$ | $2.33 \pm 0.07$ |
| $\gamma_{wat} [mN/m]^h)$ | $32^i)$ | $32 \pm 0^g)$ | $38 \pm 0$ |

a) ref <sup>90</sup> b) ref <sup>91</sup> c) ref <sup>92</sup> d) ref <sup>12</sup> e) ref <sup>93</sup> f) ref <sup>74</sup> g) Taken from the C36/LJ-PME-r simulations. h) As it is an implicit CG model (no explicit solvent molecules), it is ambiguous to distinguish surface tension at the air interface and water interface. Specific to TG, those two values are comparable (see Fig.2) i) ref <sup>75</sup>

#### TG Nucleation

TG nucleation between ER leaflets represents the first step in LD biogenesis. A previous NMR study determined the maximum solubility of TG in a PL phase to be 2.8% g TG or 2.4% mol TG.^28^ Therefore, we expected that a bilayer containing TG at a higher than critical concentration would undergo TG nucleation. Accordingly, we prepared CG bilayer systems with varying TG concentrations to study TG nucleation. Consistent with our expectations, our CG simulations confirmed that TG nucleates and forms an oil lens between the leaflets within 20M time steps when the TG concentration is 5% but does not if it is 2% (Fig. 9). In order to estimate the degree of nucleation, the nucleation % was calculated by dividing the number of TG molecules in the biggest cluster by the number of total TG molecules. If the glycerol moieties of two TG molecules are within 2 nm, they were assumed to be in the same cluster. Results for the 5% simulation indicate that the oil cluster is stable and recruits more TG with increasing simulation time until equilibrium is reached (Fig. 9). Importantly, TG molecules that are not in the oil cluster but dissolved in the membrane are primarily SURF-TG. This can be also seen in the 2% simulation where most of the TG molecules resided at the surface (Fig. 9).

**Figure 9.**
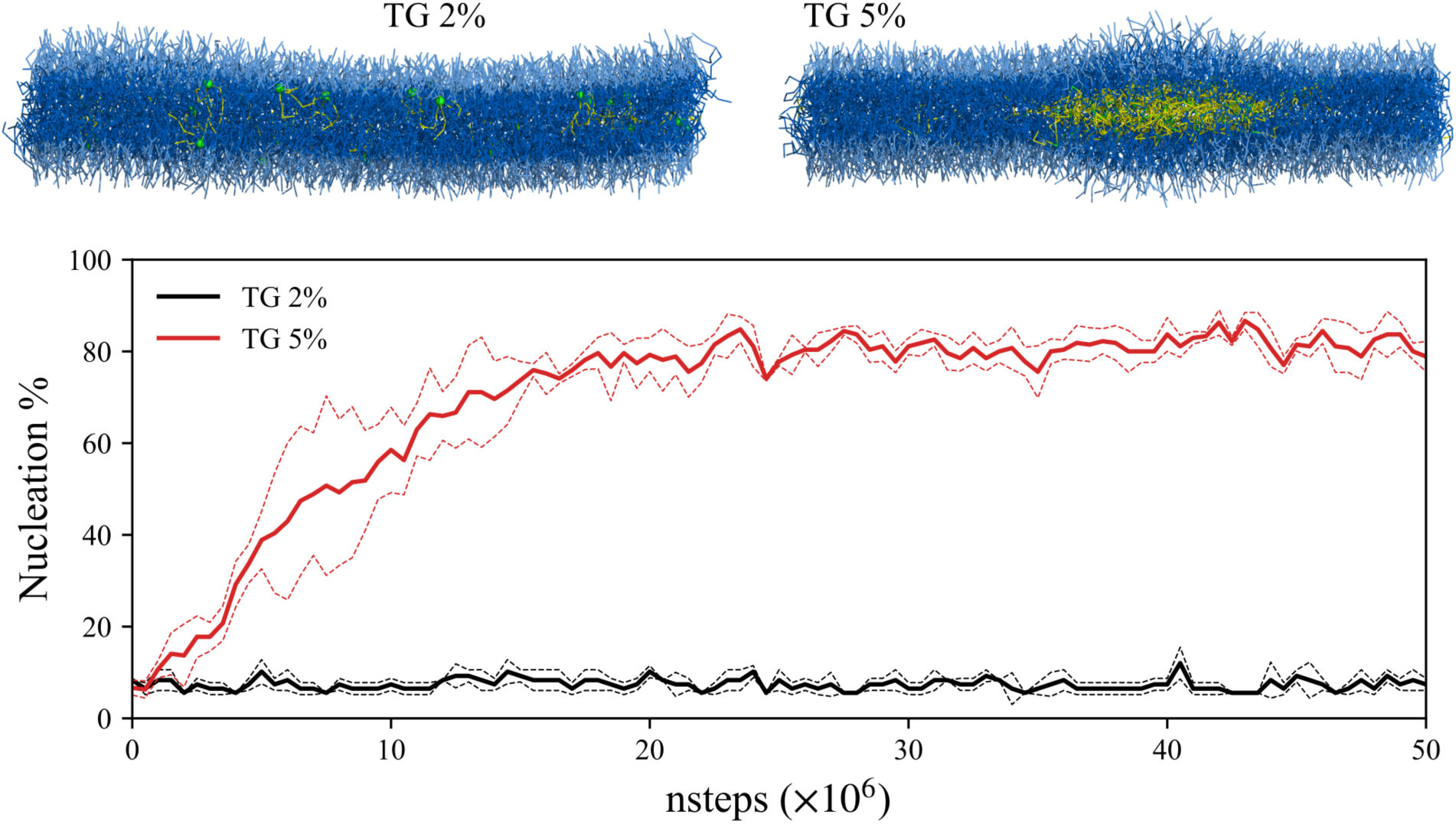
CG simulations of a symmetric bilayer and TG nucleation. The last snapshots (50M steps) of two membranes and nucleation % are shown. The dashed lines indicate the standard deviation of the quantity from three replicas for each concentration. In the snapshots, PL head group and acyl chains are shown in light blue and dark blue, respectively. The TG glycerol moiety and TG acyl chains are shown in green and yellow, respectively.

In order to show the surface propensity of a TG molecule in our force field and to evaluate the quality of the TG model, the TG flip-flop PMF was calculated with the TT-MetaD simulations (Fig. S8). Although there is a disagreement between the AA and CG results at the membrane center, the PMF suggests that TG primarily resides at the membrane surface in the absence of an oil lens.

#### Conical lipid-mediated membrane bending

As confirmed by our AA simulation results, SURF-TG locally induces a negative curvature. Using CG simulations, we investigated how local membrane deformation leads to global membrane deformation. First, we prepared an asymmetric bilayer wherein each leaflet contained a different number of SURF-TG molecules. For instance, in one of our CG simulations, the upper leaflet of the bilayer contained 3552 PL and 48 SURF-TG molecules, while the lower leaflet contained 3576 PL and 24 SURF-TG molecules. The biological rationale for the asymmetric distribution of SURF-TG between two leaflets will be discussed in the Discussion section later. Initially from a flat bilayer, the bilayer then rapidly (at 20K time steps) bent toward the lower leaflet containing less SURF-TG than the upper leaflet (Fig. 10). Interestingly, one of the regions in the upper leaflet where the local concentration of SURF-TG was initially high (x = 0.6 and y = 0.1 in Fig. 10a) became the lowest point, as indicated in Fig. 10b and Fig. 10c. The other region that initially had the high local concentration of SURF-TG in the upper leaflet (x = 0.5 and y = 0.9 in Fig. 10a) became curved to the lower leaflet. In this particular simulation, TG nucleation did not occur because of the low TG concentration. Consistent with the AA results, TG remained SURF-TG, populated at the negative curvature during 5M steps (data not shown). The same simulation but containing the higher TG concentration demonstrates the same bending behavior, followed by nucleation.

**Figure 10.**
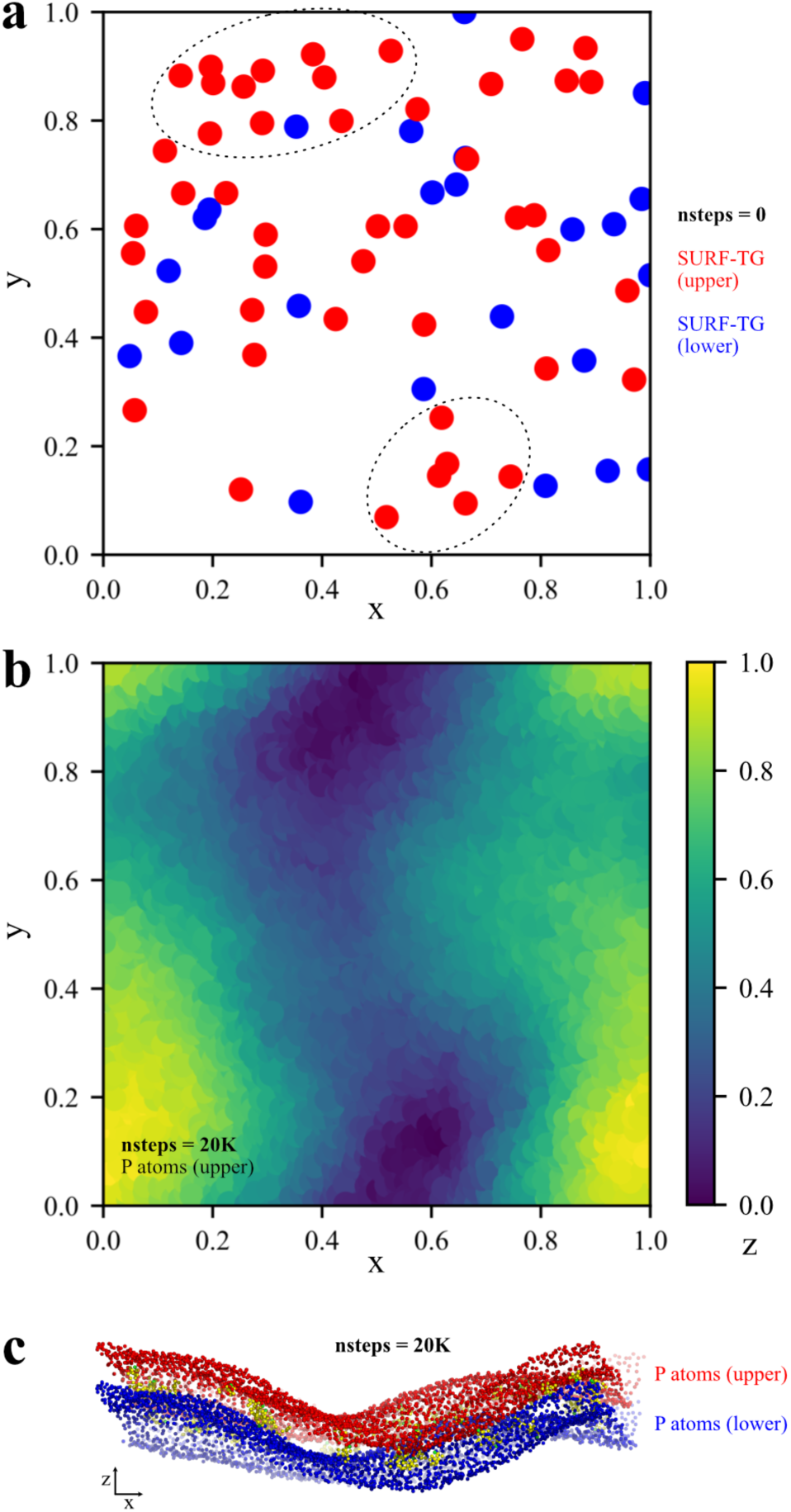
CG simulation of an asymmetric bilayer and membrane bending toward the lower leaflet. a) The initial distribution of SURF-TG in the upper leaflet (red) and in the lower leaflet (blue). The upper leaflet contains more SURF-TG than the lower leaflet; the scaled X, Y coordinates were used. The regions where the local density of SURF-TG is high in the upper leaflet are circled. b) The height of the phosphorus atoms of the upper leaflet at 20K steps. The scaled X, Y, and Z coordinates were used. c) The CG-MD snapshot at 20K time steps; the upper phosphorus atoms are shown in red and the lower phosphorus atoms in blue. TG glycerol moiety and acyl chains are shown in green and yellow, respectively.

To show the other example of conical lipid-mediated membrane bending, we prepared a bilayer immersed in water (nanodisc) and carried out CG simulation with the SDK force field (Figure 11).^68-70^ Initially, the upper leaflet contains 576 POPC molecules and the lower leaflet contains 522 POPC and 54 DOPE molecules. After 20 ns (2M MD timesteps), the membrane becomes bent toward the upper leaflet and the bent structure was maintained toward the end of the simulation (28.3 ns). One expects the nanodisc will become flat eventually once the PL distribution becomes equal.

**Figure 11.**
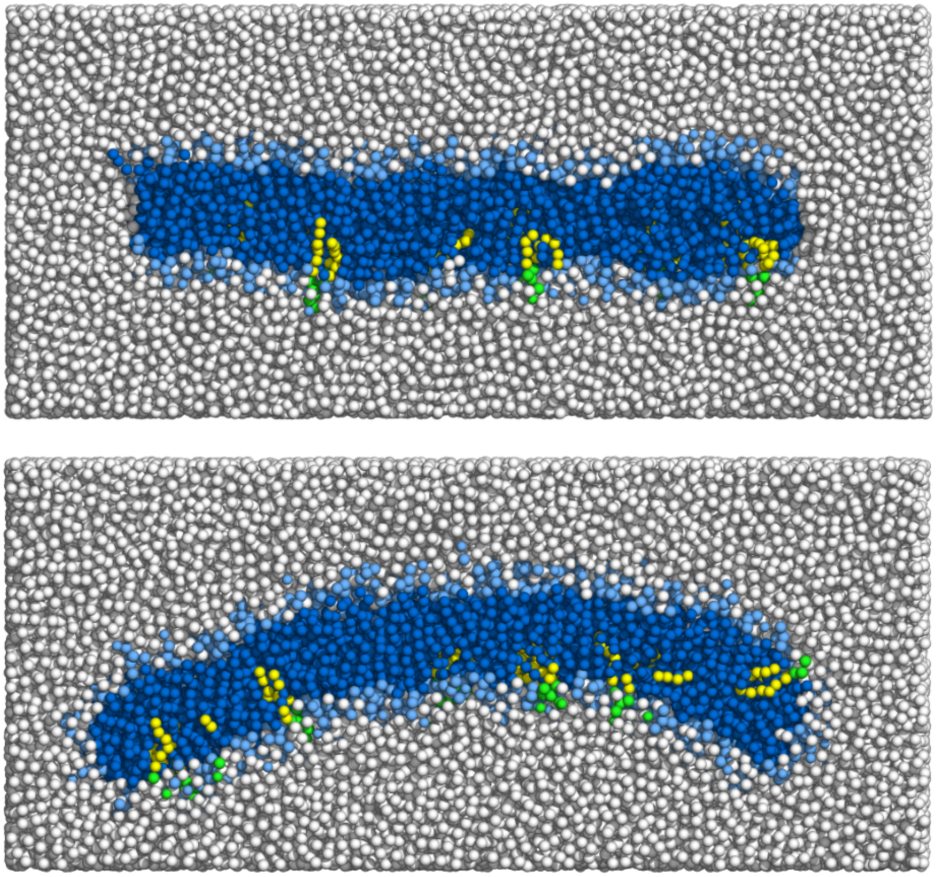
Initial (top) and 28.3 ns (bottom) snapshots of a bilayer membrane. Clipped in the XZ plane. POPC head group and tails and DOPE head group and tails are shown in light blue and dark blue and green and yellow, respectively. Waters are shown in white.

Based on the above results, we conclude that the timescale of membrane bending is faster than that of TG nucleation. In our CG simulations, membranes containing 5% TG undergo bending first, followed by TG nucleation. In the nanodisc simulation, membrane bending due to asymmetric distribution of conical lipids happen at the relatively fast timescale (20 ns). One may obtain comparable insight via the AA simulation results. In AA simulations, membrane undulation occurs within several hundred nanoseconds, thereby enabling us to evaluate the bending modulus from the undulation spectrum. In comparison, the bilayer containing 5% TG did not undergo TG nucleation within the AA timescale (microsecond). Therefore, both AA and CG simulations suggest the timescale of TG nucleation is slower than that of membrane undulation. Taken together, SURF-TG will be important in the LD biogenesis because it can modulate the membrane properties before transitioning into an oil lens. Especially, membrane deformation driven by SURF-TG, which would precede TG nucleation, may determine the LD budding directionality.

## DISCUSSION

Using a revised AA-TG model that reproduces the experimental surface tension at the water interface, we have examined herein how a conical lipid modulates membrane properties and induces membrane remodeling. The AA simulations demonstrate that SURF-TG decreases bending rigidity and increases PL ordering. Due to its conical shape, SURF-TG induces a local negative curvature. We also confirmed using biased simulations that the energy barrier for relocating SURF-TG to the bilayer center is ∼2 kcal/mol in the absence of an oil lens. This finding is consistent with our experience that TG initially locates at the bilayer center, while becoming SURF-TG within 10 ns of unbiased MD simulations and it resides principally at the surface. Finally, we systematically demonstrated the impact of neutral lipids on the membrane properties with varying DAG concentrations by determining that neutral lipids populate and induce a negative curvature and reduce the bending modulus.

In order to access SURF-TG-mediated membrane remodeling, we developed a phenomenological implicit solvent CG model for PL and TG using a Gaussian function for attraction and the LJ potential for repulsion. Primarily parameterized to reproduce the RDFs from the atomistic CG mapped trajectories, the CG model was able to reproduce most of the physical properties, with the exception of the isothermal compressibility of TG, which was found to be a factor of three higher; additionally, the bilayer bending modulus was noted to be 1.4 times higher compared with our AA results. Using our phenomenological CG model, we showed TG forms an oil lens between the leaflets when the TG concentration is 5% but does not if it is 2%. We also demonstrated that asymmetric SURF-TG composition induces membrane bending because SURF-TG works as a negative curvature inducer. For instance, when the upper leaflet of a bilayer contained more SURF-TG, the membrane would bend toward the lower direction. Consistent with the AA results, TG molecules that do not belong to the oil cluster are mostly SURF-TG.

Although a related hypothesis suggesting that asymmetric surface tension determines the directionality of LD budding has recently been proposed,^94-95^ the cause of any asymmetric surface tension was not linked to lipid type or lipid geometry. A possible physiological explanation for the asymmetric distribution of SURF-TG between the luminal leaflet and cytosolic leaflet of the ER bilayer can be found in the recently resolved structure of DGAT1.^96-97^ Catalyzing the last step of TG synthesis, DGAT1 intakes DAG and outputs TG. The lateral gate, which is located closer to the luminal leaflet, is anticipated to be a pathway for the flow of the reactant and product. Therefore, we can propose a model in which a newly synthesized TG molecule bends the membrane toward the cytosol and determines the budding directionality based on the following steps: 1) The synthesized TG molecule is released through the lateral gate of DGAT1. 2) The released TG molecule becomes SURF-TG since it has an energy penalty of ∼2 kcal/mol when residing at the ER bilayer center. 3) Given that the lateral gate is closer to the ER luminal leaflet, it is more likely that the luminal leaflet will contain more SURF-TG than the cytosolic leaflet. 4) The accumulation of SURF-TG in the luminal leaflet bends the ER bilayer toward the cytosolic side. 5) The curved ER bilayer recruits the lipid droplet assembly complex.^98^

This report makes reference to a number of papers that have described comparable results, although using different lipids and in different contexts.^93, 99-103^ In those studies, the amount of polyunsaturated fatty acids or polyunsaturated PLs were correlated with bending modulus and the distribution of those lipids. The central conclusion from those studies is that polyunsaturated acyl chains add fluidity in the Z dimension, which implies that they can be more flexibly adapted for curvature than saturated acyl chains, thereby reducing bending modulus, facilitating lipid trafficking, and modulating membrane dynamics. In this paper, TG and DAG were modeled with triolein and dioleoyl-glycerol, wherein each chain was mono-unsaturated. Therefore, we suggest that our findings are more related to the molecular shape rather than to the degree of unsaturation. Nonetheless, a future study should be designed to investigate the interplay between those two factors.

Finally, we discuss here our future work on CG modelling. In this study, a phenomenological CG model was developed via hand-tuning. A follow-on study may incorporate a bottom-up approach that systematically constructs CG models based on underlying AA interactions. Such a methodology will include the force-matching method,^104-108^ relative entropy minimization,^109^ or a hybrid approach that utilizes both methods.^110^ Using a highly coarse-grained membrane, large-scale membrane deformations mediated by the Bin/amphiphysin/Rvs (BAR) domain-containing proteins have been successfully described with such methods.^111-115^ Also, one interesting potential direction would be to introduce internal “states” to lipid CG beads using the ultra-coarse-graining (UCG) theory in order to modulate the CG interactions in different chemical or physical environments.^116-118^ In particular, by assuming that the internal state dynamics remain in quasi-equilibrium,^119-121^ we expect that the UCG modeling can further capture different TG solubilities at PL bilayers and LD surfaces.

## CONCLUSIONS

In this study we show how neutral lipids are able to modulate the physical properties of bilayers. Results from our AA-MD bilayer simulations indicate that TG principally resides at the bilayer surface, adopting PL-like conformations, in the absence of an oil lens. SURF-TG is an innately conical lipid because of the addition of the acyl chain and the elimination of the head group compared to POPC; as such, a conical SURF-TG produces a local negative curvature and lowers bending modulus. We also find a conical molecule, DAG, populates the negative curvature and lowers bending modulus. In order to increase the accessible simulation time scale, a phenomenological CG model for PL and TG was developed by parameterizing the non-bonded interactions to reproduce the RDFs from the mapped atomistic trajectories and physical properties. In the CG simulations, TG molecules form an oil cluster when TG concentration is above the critical concentration. Our CG simulations of the bilayers, wherein each monolayer surface contained a different number of SURF-TG, confirmed SURF-TG-driven membrane deformation, which may determine the directionality of LD budding and catalyze TG nucleation. Consistent with the AA results, TG molecules that are not in the oil cluster but dissolved in the PL phase are mostly SURF-TG and populate the negative curvature. To conclude, this paper demonstrates how the conical shape of neutral lipids are implicated in the early step of LD biogenesis.

## SUPPORTING INFORMATION

Description of the CG model, comparison of the 2D/3D RDF computed from the CG and CG mapped atomistic trajectories for DOPC/TG, comparison of the RDF of the TG glycerol moiety of three different TG models, PMF (C36 results) obtained with REUS and TT-MetaD, orientation and position of TG molecules in a bilayer membrane, height field and DAG distribution of the DAG 30% membrane, and TG flip-flop PMF comparison.

## ACKNOWLEDGEMENTS

This research was supported by a grant from the National Institute of General Medical Science (NIGMS) of the National Institutes of Health (NIH), grant R01-GM063796. The computer simulations were performed on the high-performance GPU cluster (GM4) at the University of Chicago Research Computing Center, supported by NSF grant DMR-1828629 and the Stampede2 supercomputer at the Texas Advanced Computing Center (TACC) through allocation TG-MCA94P017 with resources provided by the Extreme Science and Engineering Discovery Environment (XSEDE) supported by NSF grant ACI-1548562. We thank Chenghan Li, Jeeyun Chung, Zack Jarin, Myong In Oh, Jessica Swanson, Robert Farese, Jr., Tobias Walther, and Richard Pastor for their useful perspectives and discussions. We thank Stefano Vanni for providing his SDK force field of neutral lipids.

## TOC IMAGE

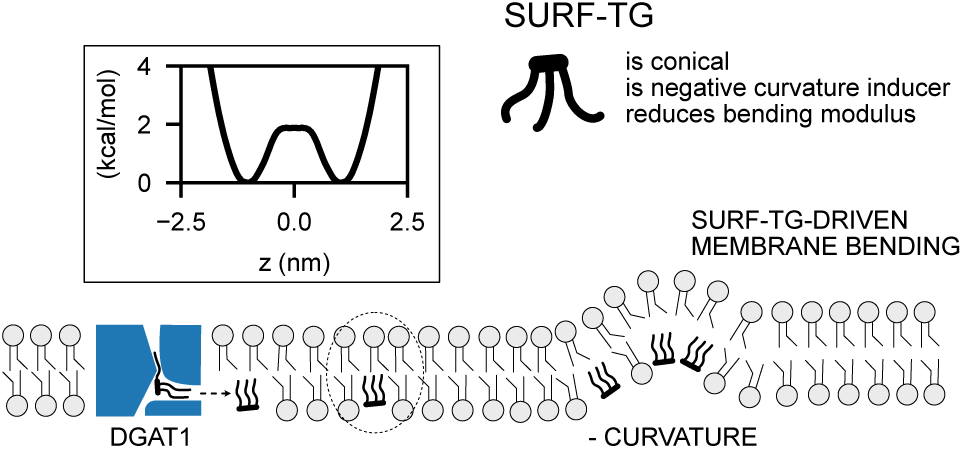

