## Supplemental File for "Physical Characterization of Triolein and Implications for Its Role in Lipid Droplet Biogenesis"

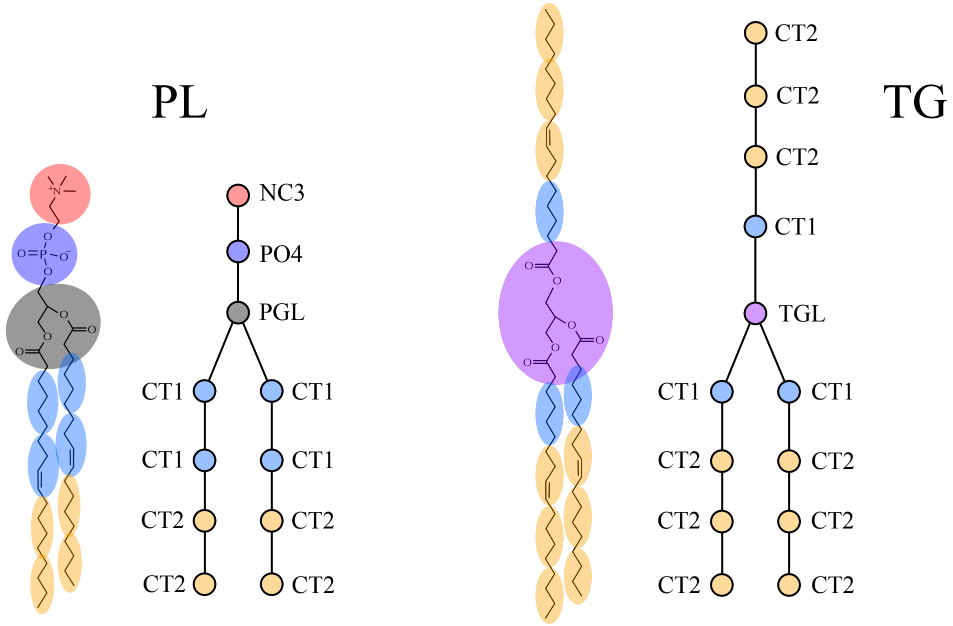


| Atom 1 | Atom 2 | A [kcal/mol] | σ [nm] |
|---|---|---|---|
| NC3 | NC3 | 0 | 0.95 |
| PO4 | PO4 | 0 | 0.70 |
| PGL | PGL | 0 | 0.75 |
| CT1 | CT1 | 1.5 | 0.68 |
| CT2 | CT2 | 0.8 | 0.69 |
| CT1 | CT2 | 1.0 | 0.685 |
| TGL | TGL | 1.5 | 0.8 |
| PGL | TGL | 1.0 | 0.775 |

**Figure S1**. Description of the CG model. The potentials used for this model are the Gaussian function as the attraction, $-A\ \exp\left(-Br^2\right)$ where $B$ was set to $2\ nm^{-2}$, and the LJ repulsive part as the repulsion, $4\varepsilon\ (\sigma/r)^{12}$ where $\varepsilon$ was set to $0.0028\ kcal/mol$. The parameters for atom $i$ and atom $j$, where $i\ \neq j$, $A_{ij} = \sqrt{A_i A_j}$ and $\sigma_{ij} = 0.5\left(\sigma_i + \sigma_j\right)$, unless otherwise specified. The mass of NC3, PO4, PGL, CT1, CT2, TGL, and PGL are approximately as-is in the mapping, which are 87, 95, 157, 55.9, 55.9, and 215 $amu$, respectively. The equilibrated bond distance and force constant are 0.5 $nm$ and 500 $kcal/mol \cdot nm^2$, respectively. The CG topology and force field are available at https://github.com/ksy141/SK_CGFF.git.

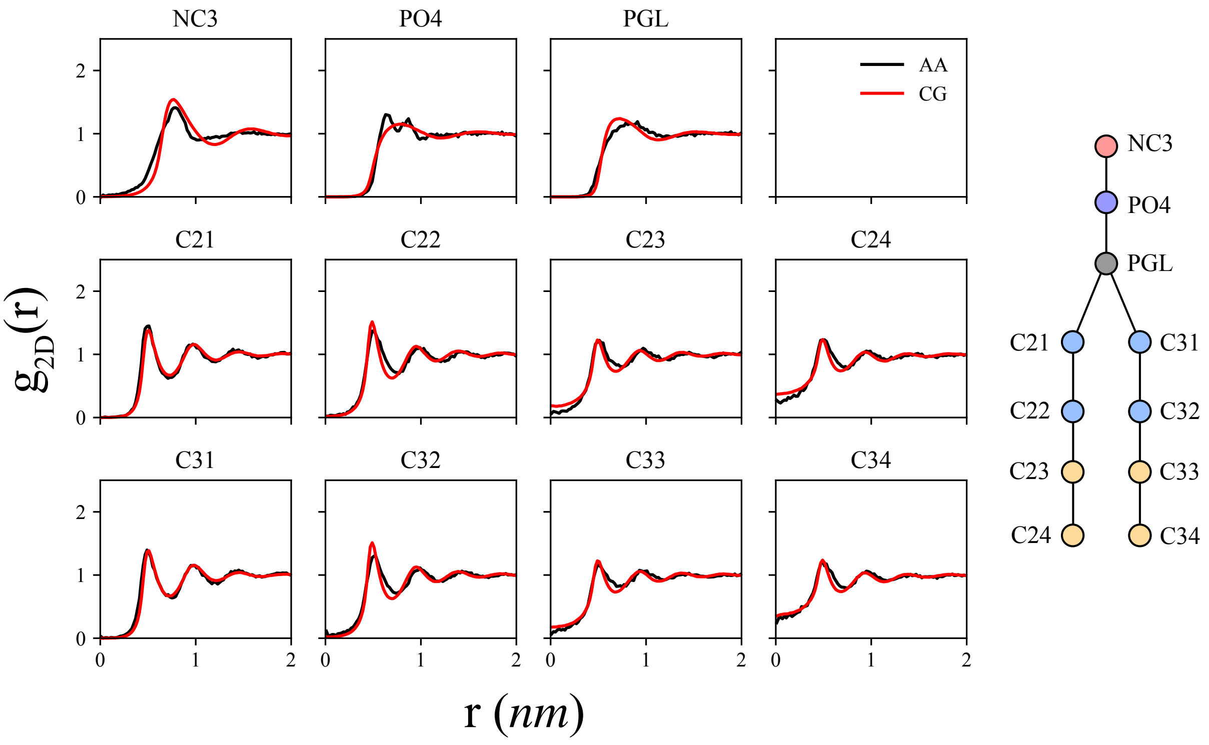


**Figure S2.** Comparison of the 2-dimensional radial distribution functions computed from the CG (red) and CG mapped atomistic (black) trajectories for DOPC. The AA simulation was carried out with C36 using a cutoff distance of 1.2 nm.

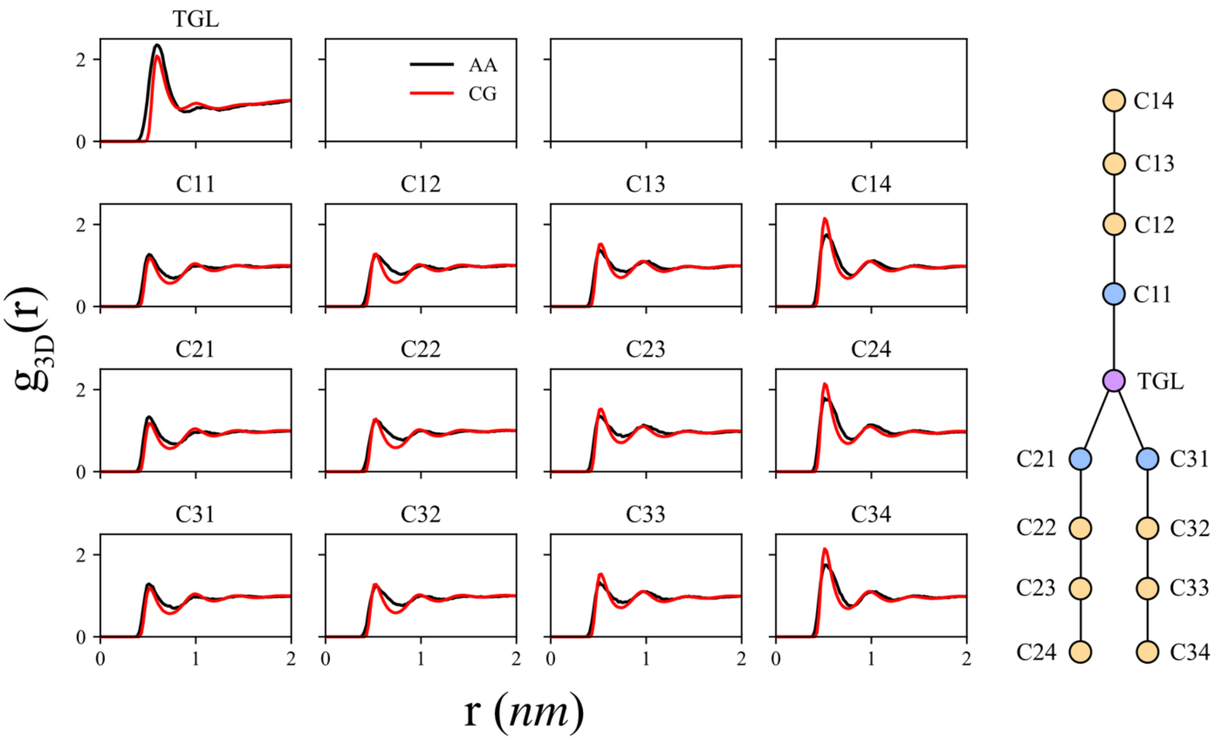


**Figure S3.** Comparison of the radial distribution functions computed from the CG (red) and CG mapped atomistic (black) trajectories for TG. The AA simulation was carried out using C36 with a cutoff distance of 1.2 nm.

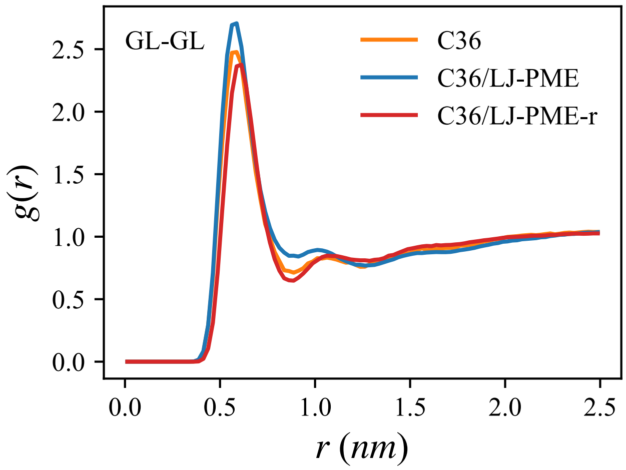


**Figure S4.** Comparison of the radial distribution functions of the TG glycerol moiety computed from CG mapped atomistic trajectories of three different TG models.

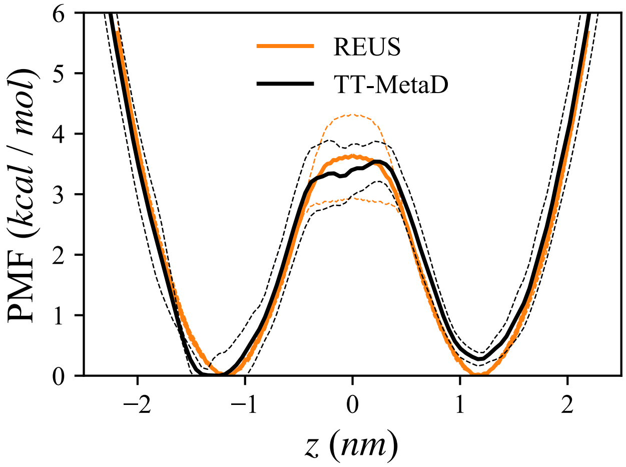


**Figure S5.** PMF obtained with REUS (orange) and TT-MetaD (black). The simulations were performed with C36 using a cutoff distance of 1.2 nm. The dashed lines indicate the standard deviation. Five equal-length blocks and five replicas were used for REUS and TT-MetaD, respectively, to estimate the standard deviation. The REUS PMF was reflected with respect to the membrane center. Related to Fig. 3 of the main text.

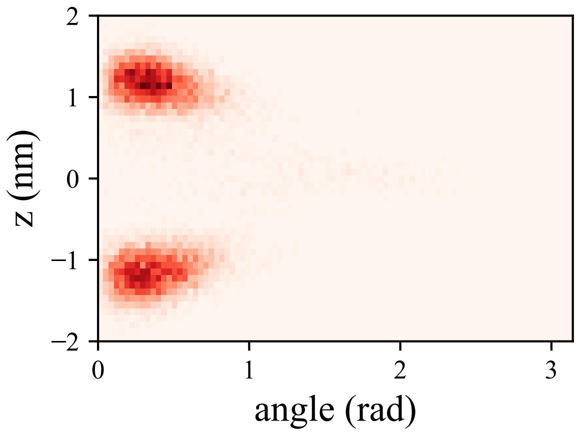


**Figure S6.** Orientation and position of TG molecules in a bilayer membrane. The Z position of 0 represents the membrane center. The angle is between the Z-axis and the positional vector of the TG glycerol moiety from the center of the mass of the TG acyl chains.

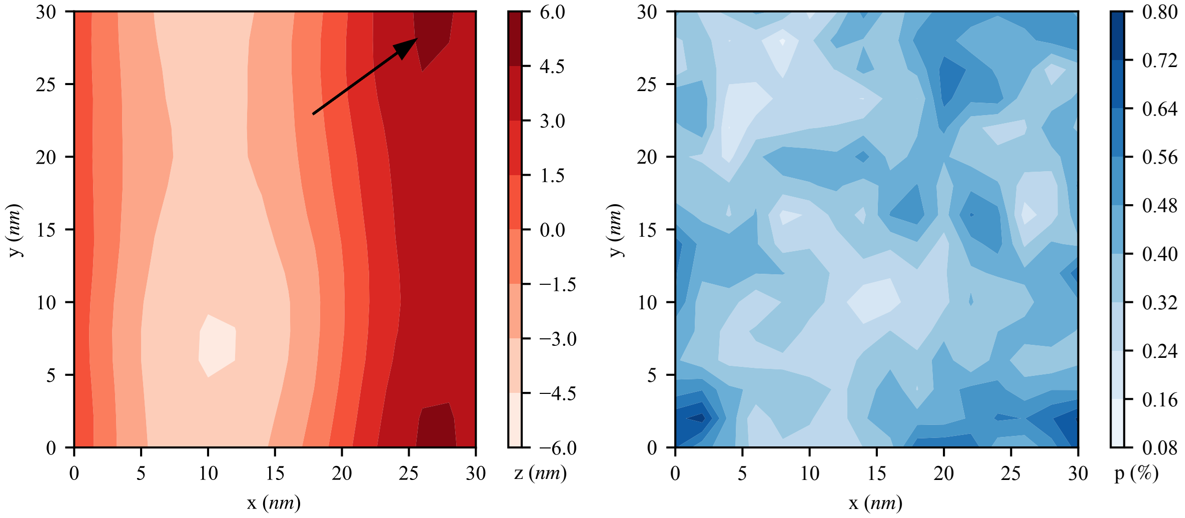


**Figure S7.** Height field (left) and DAG distribution (right) of the lower leaflet of the DAG 30% membrane. An arrow in the left figure indicates an exemplary motion of a DAG molecule diffusing into the negative curvature region during 500 ns. The total length of simulation was 1 µs. Related to Fig. 8 of the main text.

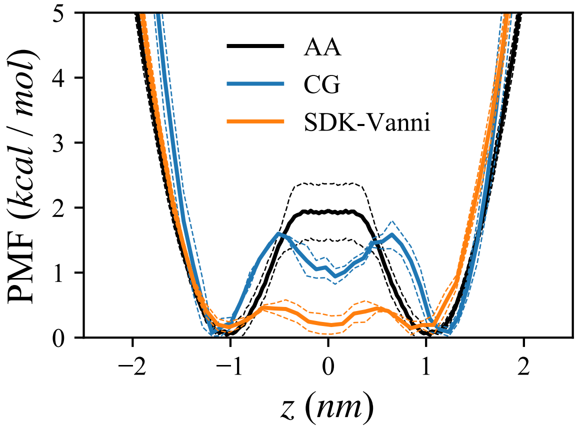


**Figure S8.** TG flip-flop PMF comparison, calculated with the C36/LJ-PME-r force field (black), the CG force field developed in this manuscript (blue), and the Vanni's modified SDK force field (orange). In the CG simulations, three replicas were used to estimate the standard deviation, shown in the dashed lines.
